# Exchange of a single amino acid residue in the cryptophyte phycobiliprotein lyase *Gt*CPES expands its substrate specificity

**DOI:** 10.1101/2020.03.31.018853

**Authors:** Natascha Tomazic, Kristina E. Overkamp, Marco Aras, Antonio J. Pierik, Eckhard Hofmann, Nicole Frankenberg-Dinkel

## Abstract

Cryptophyte algae are among the few eukaryotes that employ phycobiliproteins (PBP) for light harvesting during oxygenic photosynthesis. In contrast to the cyanobacterial PBP that are organized in large membrane-associated super complexes, the phycobilisomes, those from cryptophytes are soluble within the chloroplast thylakoid lumen. Their light-harvesting capacity is due to covalent linkage of several open-chain tetrapyrrole chromophores (phycobilins). *Guillardia theta* utilizes the PBP phycoerythrin PE545 with 15,16-dihydrobiliverdin (DHBV) in addition to phycoerythrobilin (PEB) as chromophores. Thus far, the assembly of cryptophyte PBPs is not yet completely understood but involves the action of PBP-lyases as shown for cyanobacterial PBP. PBP-lyases facilitate the attachment of the chromophore in the right configuration and stereochemistry. Here we present the functional characterization of eukaryotic S-type PBP lyase *Gt*CPES from *G. theta*. We show *Gt*CPES mediated transfer and covalent attachment of PEB to the conserved Cys^82^ of the acceptor PBP β-subunit (*Pm*CpeB) of *Prochlorococcus marinus* MED4. Based on the previously solved crystal structure, the *Gt*CPES binding pocket was investigated using site-directed mutagenesis. Thereby, amino acid residues involved in phycobilin binding and transfer were identified. Interestingly, exchange of a single amino acid residue Met^67^ to Ala extended the substrate specificity to phycocyanobilin (PCB) likely by enlarging the substrate-binding pocket. Variant *Gt*CPES_M67A binds both PEB and PCB forming a stable, colorful complex *in vitro* and *in vivo* produced in *Escherichia coli*. *Gt*CPES_M67A is able to mediate PCB transfer to Cys^82^ of *Pm*CpeB. Based on our data we postulate that a single amino acid residue determines the bilin-specificity of phycoerythrin S-type lyases but that additional factors regulate hand over to the target protein.

---

Phycobiliproteins (PBPs) are the photosynthetic light-harvesting structures of cyanobacteria, red algae and cryptophytes and are used to absorb regions of the visible light spectrum that are poorly covered by chlorophylls. Consequently, PBPs efficiently absorb light in the spectral region between 430 and 680 nm (1) They consist of apo-proteins with covalently linked open-chain tetrapyrrole molecules termed phycobilins. While PBPs in cyanobacteria and red algae are organized in large light-harvesting structures called phycobilisomes, those of cryptophyte algae are soluble in the thylakoid lumen of the chloroplast (2–4). Cryptophycean PBPs occur in large concentrations and are organized in (αβ)(α’β)-heterodimers (4). In contrast to cyanobacteria, cryptophytes only use a single type of PBP. The model species *Guillardia theta* utilizes phycoerythrin 545 (PE545) with an absorption maximum at 545 nm (5). It consists of the α-subunit CPEA with bound 15,16-dihydrobiliverdin (DHBV) and the β-subunit CpeB loaded with three molecules of phycoerythrobilin (PEB) (6,7). The *G. theta* nuclear genome encodes 16 CPEA copies that share high homology among each other but lack homologies to any other known protein (8). In contrast, the plastid-encoded β-subunit is highly similar to those of cyanobacteria and rhodophytes (2,9–12). Both subunits possess thioether-linked phycobilins at conserved cysteine residues (α-Cys^19^; β-Cys^82^, β-Cys^155^), one is connected via a double linkage (β-Cys^50/61^) (4). The biosynthesis of the open-chain tetrapyrrole chromophores is highly conserved among PBP containing organisms, and starts with cleavage of a heme molecule by heme oxygenase. The resulting product biliverdin IXα (BV) is the substrate of enzymes belonging to the ferredoxin-dependent bilin reductases and reduce the BV substrate at different positions (13,14). *G. theta* was shown to possess a plastid encoded heme oxygenase and two nucleus encoded FDBRs, PEBA and PEBB involved in the conversion of BV to PEB via 15,16-DHBV (14). Following their synthesis, the phycobilins are post-translationally attached to the PBP with the help of PBP-lyases, which mediate the efficient attachment to the specific cysteine residues of the apo-PBPs (15).

PBP-lyases are classified into E/F-, S/U- and T-type lyases, each being responsible to facilitate the attachment of a phycobilin to a specific cysteine residue (15) Within the genomes of *G. theta* we previously identified four different PBP-lyases: CPES, CPET, CPEX and CPEZ. All are encoded in the nuclear genome, except CPET, which was identified on the nucleomorph genome (14,16). *Gt*CPES from *G. theta* belongs to the S/U-type lyases and adopts a 10-stranded β-barrel with a modified lipocalin fold (14). It specifically binds phycobilins with a reduced C15,C16-bond like 15, 16-DHBV and PEB (14). Binding of phycocyanobilin (PCB) with a rigid double bond between C15 and C16 was not observed. Based on a study of a CPES homolog from the cyanobacterium *Prochlorococcus marinus* MED4, *Gt*CPES is hypothesized to be involved in the attachment of PEB to β-Cys^82^ (17). Likely, the lyase is not catalyzing the transfer reaction itself, but rather assists in the handover of the phycobilin in the correct configuration to allow stereospecific autocatalytic attachment.

In this study, we demonstrate the specific ability of *Gt*CPES to support the transfer of PEB to Cys^82^ of CpeB from the cyanobacterium *Prochlorococcus marinus* MED4 (*Pm*CpeB). Moreover, we identified individual amino acid residues involved in phycobilin binding and transfer with one single amino acid change expanding the substrate spectrum. Taken together our data emphasize that the narrow substrate specificity of the PE specific S-type PBP-lyases are due to a single amino acid residue.

## Results

### GtCPES mediates transfer of PEB to Cys^82^ of PmCpeB

Although we have previously shown that *Gt*CPES binds PEB and 15,16-DHBV with high affinity, an involvement in the chromophorylation of the *G. theta* PBP β-subunit at β-Cys^82^ has not yet been shown. This was mainly due to solubility issues of the recombinant *G. theta* CpeB apo-protein and we were not able to solve this issue until now (14). Furthermore, it is still unclear whether chromophorylation of PBP has to follow a specific order. In this regard, it has previously been reported that β-Cys^155^ is the first site of chromophorylation in phycocyanin (18) Therefore, we took another approach to investigate whether *Gt*CPES is able to support PEB attachment to the PE β-subunit. We made use of β-PE (CpeB) from the cyanobacterium *P. marinus* MED4 (*Pm*CpeB). This β-subunit is the only PBP in this organism and is highly degenerated, as it only possesses a single chromophorylation site at Cys^82^ (19,20). Recombinant *Pm*CpeB can be purified in adequate amounts to test chromophore transfer employing *Gt*CPES (17). To verify whether *Gt*CPES is sufficient for correct bilin addition to *Pm*CpeB we compared spontaneous and lyase mediated attachment of *3*(*Z*)-PEB to *Pm*CpeB by recording the fluorescence emission of assembled holo-CpeB. Covalent attachment was furthermore confirmed by zinc-blot analysis. *Pm*CpeB lacking the cysteine attachment site (i.e. *Pm*CpeB_C82A) served as a negative control (Figure 1). In all experiments, the lyase was used in excess to prevent the presence of free bilin in the sample. Spontaneous and lyase-mediated attachment resulted in the formation of covalent fluorescent phycobiliprotein complexes (Figure 1AB). Reaction products (i.e. holo-*Pm*CpeB) were stronger and displayed a more stable fluorescence in presence of *Gt*CPES with an emission maximum at 568 nm (λ_ex_=550 nm). Addition of free 3(*Z*)-PEB to apo-*Pm*CpeB resembled spontaneous attachment as described before (17). These measurements are in agreement with data observed for the native *Prochlorococcus* lyase CpeS (17) and confirm the activity of *Gt*CPES in mediating the correct attachment of PEB to β-Cys^82^. Interestingly, when we employed an *in vivo* assembled Gt*CPES*:PEB complex (obtained by coexpressing *Gt*CPES with PEB biosynthesis genes), we obtained a fluorescent product that resembled even better the native holo-*Pm*CpeB with a fluorescence emission at 571 nm (data not shown).

**FIGURE 1.**
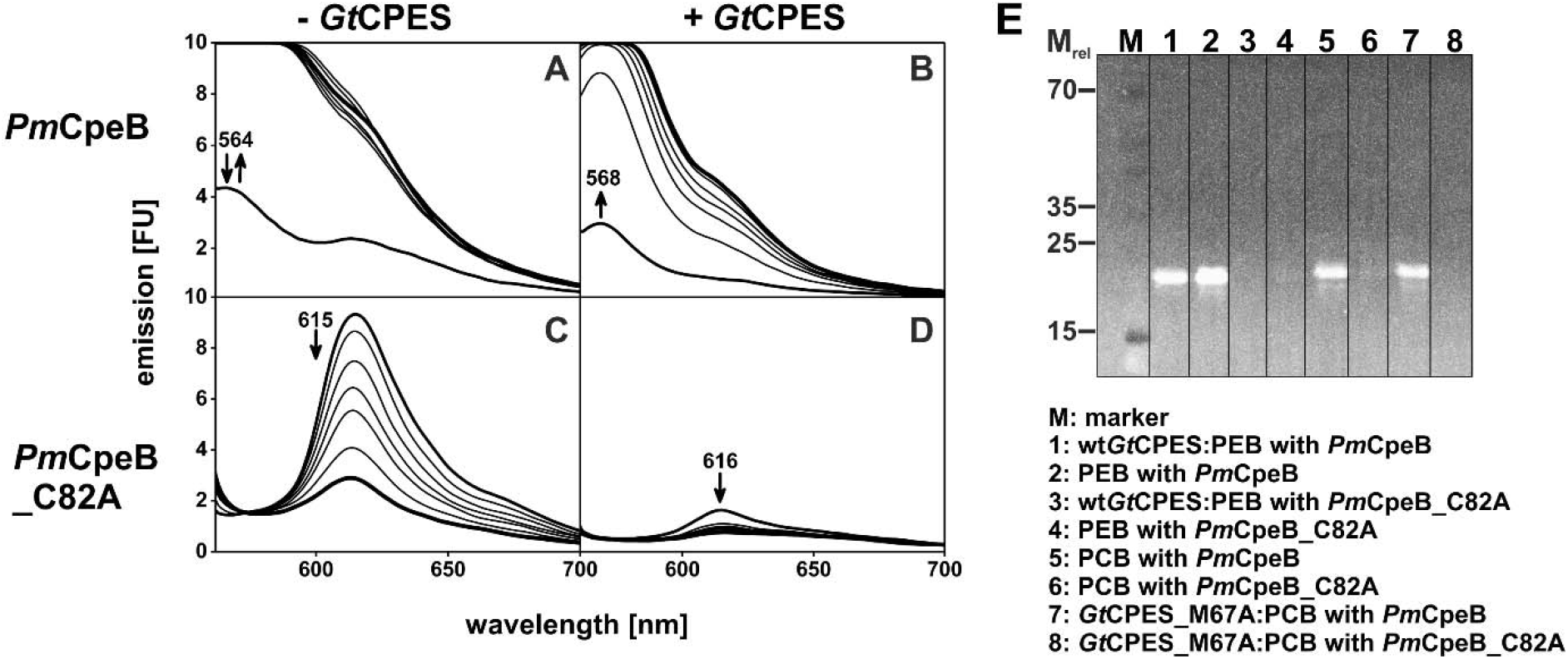
*Gt*CPES mediated and spontaneous PEB chromophorylation of *Pm*CpeB or *Pm*CpeB_C82A and covalent binding formation of *Pm*CpeB with PCB or PEB visualized by zinc-enhanced fluorescence. The apo-PBPs *Pm*CpeB or *Pm*CpeB_C82A were incubated with PEB (**A**, **C**) or wt*Gt*CPES:PEB complexes (**B**, **D**). Emission spectra (λ_ex_ = 550 nm) of 1, 5, 10, 15, 20, 30, and 45 min after addition of *Pm*CpeB or *Pm*CpeB_C82A, respectively, were detected. First and last spectra are shown by bold lines. Emission maxima are given and the course of fluorescence emission changes is indicated by arrows. After spontaneous and PBP-lyase mediated transfer studies of PEB and PCB, samples were used for SDS-PAGE and blotted onto PVDF membrane (**E**). Membrane was incubated for 1 h at 4 °C in 1.3 M zinc acetate and detected under UV-light (312 nm). Fluorescence signals based on zinc-phycobilin complexation describe covalent binding of phycobilins to *Pm*CpeB (with his-tag: 21.4 kDa). Absent signals indicate missing binding.

Our negative control, *Pm*CpeB_C82A, displayed basal interaction with free 3(*Z*)-PEB visualized by an initial increase but subsequent decrease in fluorescence emission at 615 nm (λ_ex_=550 nm) (Figure 1C). Employing the PEB-loaded *Gt*CPES lyase, no transfer was observed indicated by negligible fluorescence emission at 616 nm (λ_ex_=550 nm) (Figure 1D). As expected, no covalent attachment was observed in zinc-blot analysis (Figure 1E). Although we have previously shown that *Gt*CPES can also bind the semi reduced intermediate 15,16-DHBV (14), DHBV-loaded lyase did not result in transfer of DHBV to *Pm*CpeB (data not shown). As previously shown, DHBV is also not able to spontaneously attach to *Pm*CpeB (17).

### Two glutamate residues are important for PEB binding

Now that we established the assistance (chaperon) function of *Gt*CPES during PEB transfer, we went ahead to perform a rigorous analysis to identify amino acid residues involved in substrate binding and transfer. Despite several approaches for co-crystallization of the lyase with its substrate PEB, no co-crystal structure was obtained. However, one molecule of 1,6-hexanediol was found within the barrel in the original structure giving a hint for the putative substrate binding site and allowing the initial identification of two residues involved in substrate binding (14). We now used the crystal structure and an amino acid sequence alignment of S-type lyases to identify ten individual amino acid residues potentially involved in PEB binding and transfer (Figure 2, Figure S1, (14)). These residues were exchanged to alanine residues using site directed mutagenesis. The following variants were generated: R18A, H21A, N22A, W69A, W75A, E136A, R146A, R148A, S150A and E168A (Figure 2). Residues E136, R146, and R148 are highly conserved among S-type lyases and are in close proximity to the 1,6-hexanediol molecule within the crystal structure of *Gt*CPES and likely involved in substrate binding/coordination (14). Most of the other amino acid residues are located within the barrel and have potential influence on PEB binding. W75, which is located at the top of the lyase barrel, almost like a lid over the barrel, has putative influence on the interaction with the apo-protein CpeB (Figure 2). All variants were produced and purified in sufficient amounts and tested for their ability to bind PEB. In addition, thermal shift assays were performed to evaluate the effects the sequence variations on protein integrity and stability (21)

**FIGURE 2.**
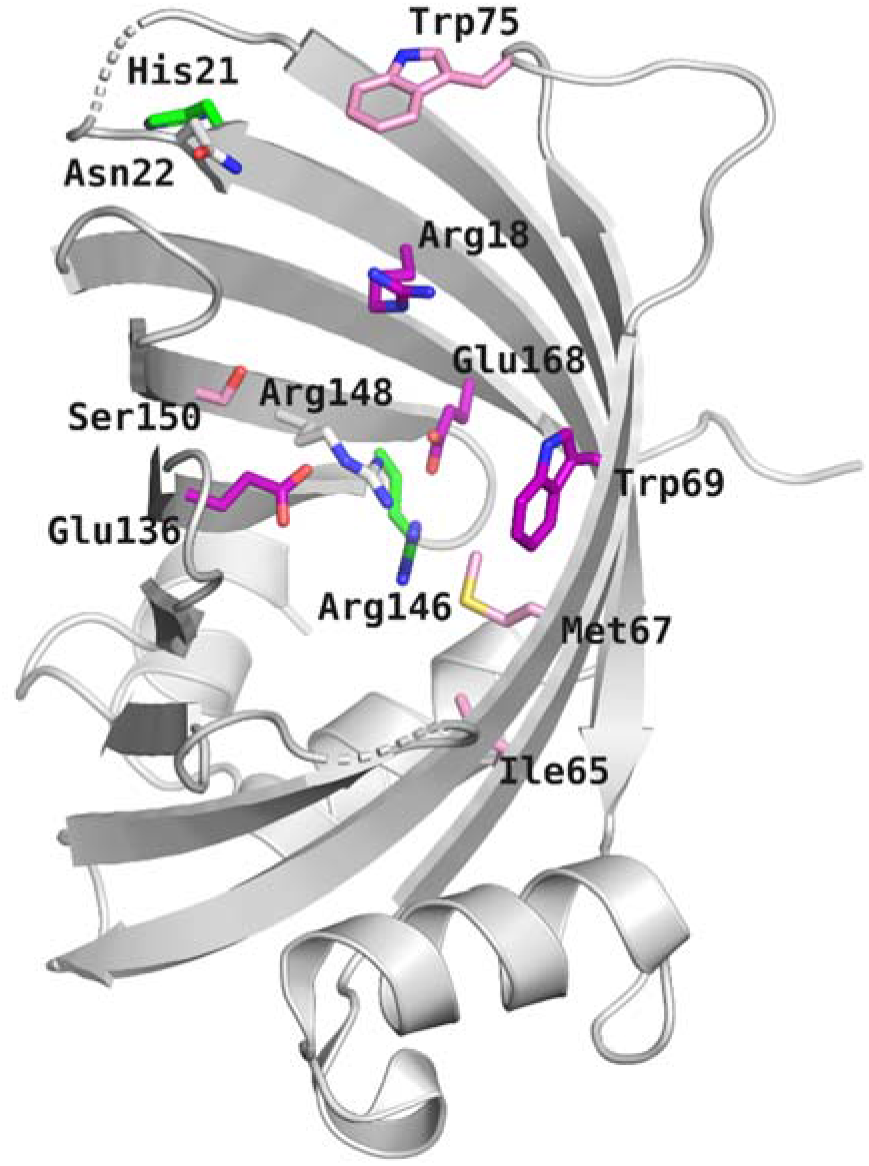
Location of amino acid residues of *Gt*CPES investigated within this study. The structure of *Gt*CPES (PDB code 4TQ2) is shown in cartoon representation. Residues discussed in the main text are shown as sticks with color coding representing the effect of a site specific exchange on *Gt*CPES function: white – no change; green – structural defect; purple – bilin binding; pink – bilin transfer and selectivity. Figure generated with Pymol (35).

Upon binding to the wt *Gt*CPES lyase, the absorbance maximum of PEB is red shifted from 535 nm to 598 nm with a significant increase in intensity with a shoulder at 557 nm (14). Five of the investigated *Gt*CPES variants (R18A, N22A, W75A, R148A and S150A) showed almost identical binding spectra with a long wavelength major peak and a shoulder at a shorter wavelength (Table 1). These variants were considered wt-like. The remaining five variants displayed moderate to considerable changes from the wt *Gt*CPES:PEB spectrum (Figure 3). The spectrum of variant W69A together with PEB still displayed two absorption maxima. However, what appears as a shoulder in the spectrum of the wt lyase, is a more distinct peak at 549 nm in this variant (Figure 3). The variants H21A and E168A also displayed two absorption peaks when incubated with PEB. However, there appears to be a large proportion of unbound PEB in these samples due to the absence of an increase in extinction coefficient of PEB upon incubation with the variants. In addition, the peak maxima were rather broad suggesting that the chromophore is less stretched, again suggesting a larger proportion of free PEB in the sample (22). In contrast, variants E136A and R146A did not bind PEB as a spectrum almost identical to that of free PEB was observed (Figure 3). The lack of PEB binding of the R146A variant is likely due to (partly) misfolded/aggregated protein. No discrete melting point between 47 and 58 °C was obtained, as contrary to other mutants. The latter was also true for variant H21A. Therefore, both amino acid residues are likely important for structural integrity of the protein. Both variants were not investigated further. Interestingly, all variants that showed changes in their absorption properties with bound PEB, also displayed fluorescence emission at ~638 nm when excited at 550 nm (Table 1). This is likely due to unspecific interaction of PEB with the lyase variants and differs from the wt protein where no fluorescence upon PEB binding was observed. In conclusion, both glutamate residues, E136 and E168 are strongly involved in PEB binding, while W69 has only minor influence on binding the substrate. H21 and R146 are important for structural integrity.

**FIGURE 3.**
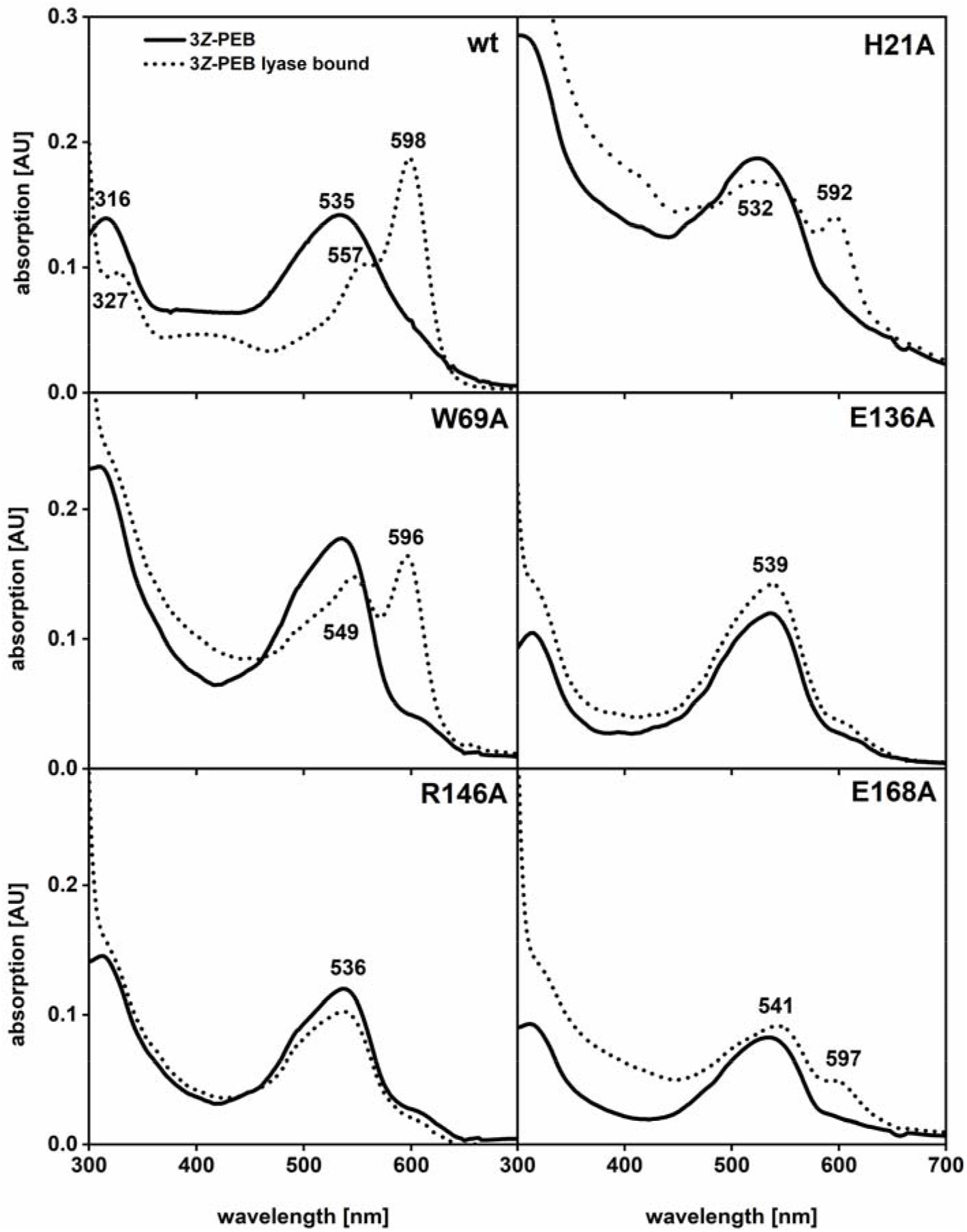
UV-Vis spectra of 3(*Z*)-PEB bound to *Gt*CPES and variants. Free PEB spectra are shown as solid lines, the spectra of lyase variants with PEB are shown as dotted lines. Peak maxima are given and the respective amino acid exchange.

**TABLE 1.** Binding and transfer of 3(*Z*)-PEB after incubation with different *Gt*CPES variants compared to the wild type. Melting points (in °C) of proteins determined in triplicate and absorbance/emission maxima of binding/transfer are given. Complex E_max_: detected emission maximum after adding PBP-lyase to free PEB. E_max_: emission maximum after 45 min incubation time of *Gt*CPES (variant):bilin with *Pm*CpeB.Excitation wavelength was 550 nm.

| *Gt*CPES<br>(variant) | $T_m$ [°C] | Binding<br>PEB | $A_{\max}$<br>[nm] | Complex<br>$E_{\max}$ [nm] | Transfer<br>PEB | $E_{\max}$<br>[nm] |
| --- | --- | --- | --- | --- | --- | --- |
| WT | 56.0 ± 0.0 | yes | 554, 600 | --- | yes | 568 |
| R18A | 48.0 ± 0.0 | yes | 552, 593 | --- | (yes)<sup>a</sup> | 567 |
| N22A | 55.0 ± 0.0 | yes | 555, 598 | --- | yes<sup>a</sup> | 568 |
| W69A | 48.0 ± 0.0 | yes | 549, 596 | 637 | no | 565 |
| W75A | 58.0 ± 0.0 | yes | 554, 599 | --- | no | n.d. |
| E136A | 47.5 ± 0.0 | no | 539 | 638 | no | 565 |
| R148A | 56.0 ± 0.0 | yes | 552, 600 | --- | yes | 566 |
| S150A | 48.2 ± 0.3 | yes | 554, 600 | --- | (yes)<sup>a</sup> | 568 |
|  | 57.5 ± 0.0 |  |  |  |  |  |
| E168A | 49.0 ± 0.0 | (no)<sup>b</sup> | 541, 597 | 642 | (no) | 568 |

<sup>a</sup> Less increase of fluorescence emission compared to wt*Gt*CPES. <sup>b</sup> Binding spectra differs from wt*Gt*CPES:PEB spectra. n.d., not detectable

### A tryptophan residue is important for PEB transfer to PmCpeB

We next tested whether the generated *Gt*CPES variants that still showed PEB binding were also able to transfer PEB to apo-*Pm*CpeB. Transfer of PEB was monitored using fluorescence spectroscopy as described above. Most of the variants that did show wt-like binding of the substrate PEB, did also display this behavior for the transfer reaction (Figure 4, S2). Interestingly, variant W69A that only showed minor absorption changes upon PEB binding (see paragraph before) did show abnormal transfer behavior that resembled that of a non-lyases mediated spontaneous reaction. Such reactions are indicative of initial increase but subsequent decrease of fluorescence emission at 565 nm with an additional fluorescence emission maxima observed at longer wavelength. Therefore, W69 likely is involved in PEB binding and its exchange leads to a large proportion of unbound PEB, which is then transferred in a spontaneous, unspecific reaction to *Pm*CpeB. The same was observed for variant E168A that already showed only weak to negligible PEB binding. This variant already displayed fluorescence emission when incubated with PEB (E_max_= 642 nm) which decreased upon incubation with *Pm*CpeB. Transfer is observed but based on the fluorescence emission, which increased and decreased, it appears that there is a large proportion of spontaneous transfer taking place. Variant S150A showed a normal PEB binding spectrum but reduced transfer indicative of a low fluorescence intensity increase upon incubation with *Pm*CpeB. The most important residue for PEB transfer is a tryptophan located at the upper rim of the barrel. Variant W75A, although still being able to bind PEB like the wt lyase, is not able to facilitate the transfer of PEB to *Pm*CpeB. In order to test whether tight binding and therefore reduced release rates of PEB might cause this observation; binding affinities of PEB to wt *Gt*CPES and the W75A variant were determined. Interestingly, *Gt*CPES_W75A is able to bind PEB more than 10-fold tighter than the wt protein suggesting that this tight binding might prevent the release and transfer of PEB (Table 2).

**FIGURE 4.**
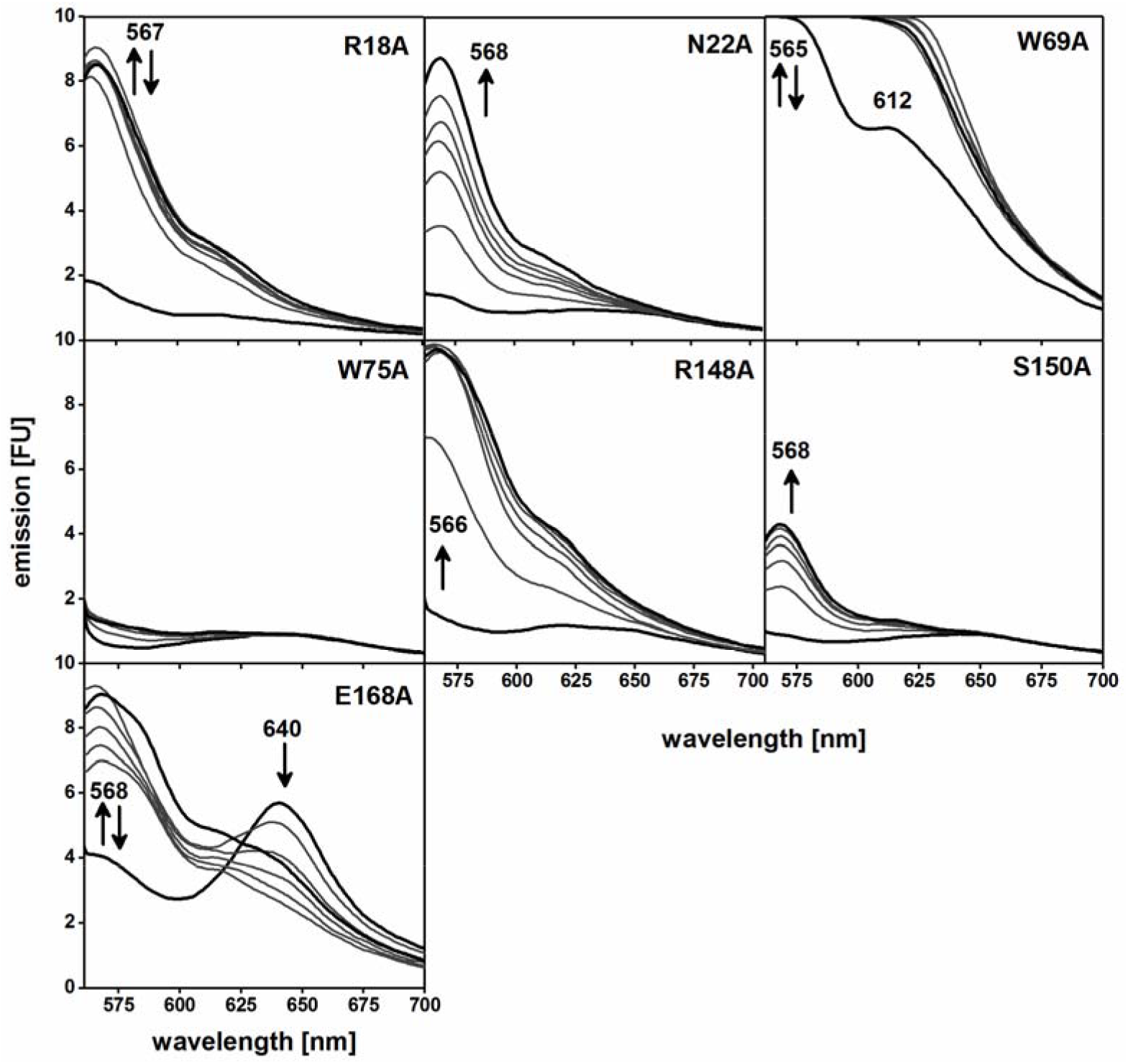
Transfer of 3(*Z*)-PEB to *Pm*CpeB by *Gt*CPES variants. Apo-*Pm*CpeB was incubated with *Gt*CPES variants preincubated with PEB and. Emission spectra (λ_ex_ = 550 nm) at 1, 5, 10, 15, 20, 30, and 45 min after addition of *Pm*CpeB were taken. First and last spectra are shown by bold lines. Emission maxima are given and the course of fluorescence emission changes is indicated by arrows.

**TABLE 2.** Binding affinities for complex formation with PEB and PCB. Phycobilin binding analysis was performed as described were tested by titration described in the experimental procedures. K_D_ values were calculated from equation (1) by using *solver* function of *Microsoft Excel*. Results were averaged over two measurements. n.b., no binding.

| *Gt*CPES<br>(variant) | PEB<br>$K_D$ [μM] | PCB<br>$K_D$ [μM] |
| --- | --- | --- |
| WT | 14.2 ± 2.6 | n.b. |
| M67A | 0.044 ± 0.003 | 7.0 ± 0.4 |
| W75A | 0.86 ± 0.06 | --- |

### A single methionine residue determines substrate specificity

*Gt*CPES possesses a narrow binding pocket harboring amino acid residues with bulky side chains. As a consequence, *Gt*CPES has a very high substrate specificity towards PEB. Nevertheless it is also able to bind the biosynthetic precursor 15,16-DHBV (14). Binding of the PEB-isomer phycocyanobilin (PCB) was not observed likely due to a rigid double bond between C15-C16 resulting in a missing flexibility of D-ring to fit into the binding pocket (Figure 5C). In order to characterize the molecular determinants of the substrate specificity, we compared both binding pocket and sequence of *Gt*CPES with the PCB-specific S-type lyase CpcS from *Thermosynechococcus elongatus* (*Te*CpcS) (Figure 5A, S1). Based on a comparison of the two available crystal structures of *Te*CpcS and *Gt*CPES, it became obvious that *Te*CpcS possesses a binding pocket with amino acid residues with less bulky side chains (14,23) We therefore went ahead to mutagenize the binding pocket of *Gt*CPES with the aim to widen it. Subsequently, the variant proteins were tested for their ability to bind PCB as well. We started to exchange amino acid residues around position 67 because these residues specifically constrict the binding pocket (Figures 2, 5). Overall, methionine 67 indeed seems to be highly specific for determining the substrate specificity. M67A and M67V variants of *Gt*CPES were still able to bind PEB as indicated by binding spectra of PEB that resembled the wt situation (Table 3). In addition, we tested whether these variants were still able to transfer PEB to *Pm*CpeB. Both methionine variants and the I64A variant displayed specific transfer of 3(Z)-PEB (Figure 6). Interestingly, the M67A variant had the strongest affinity towards PEB of all measured *Gt*CPES variants (Table 2) indicating that the affinity alone does not determine whether the chromophore is released from the binding pocket of the lyase or not.

**FIGURE 5.**
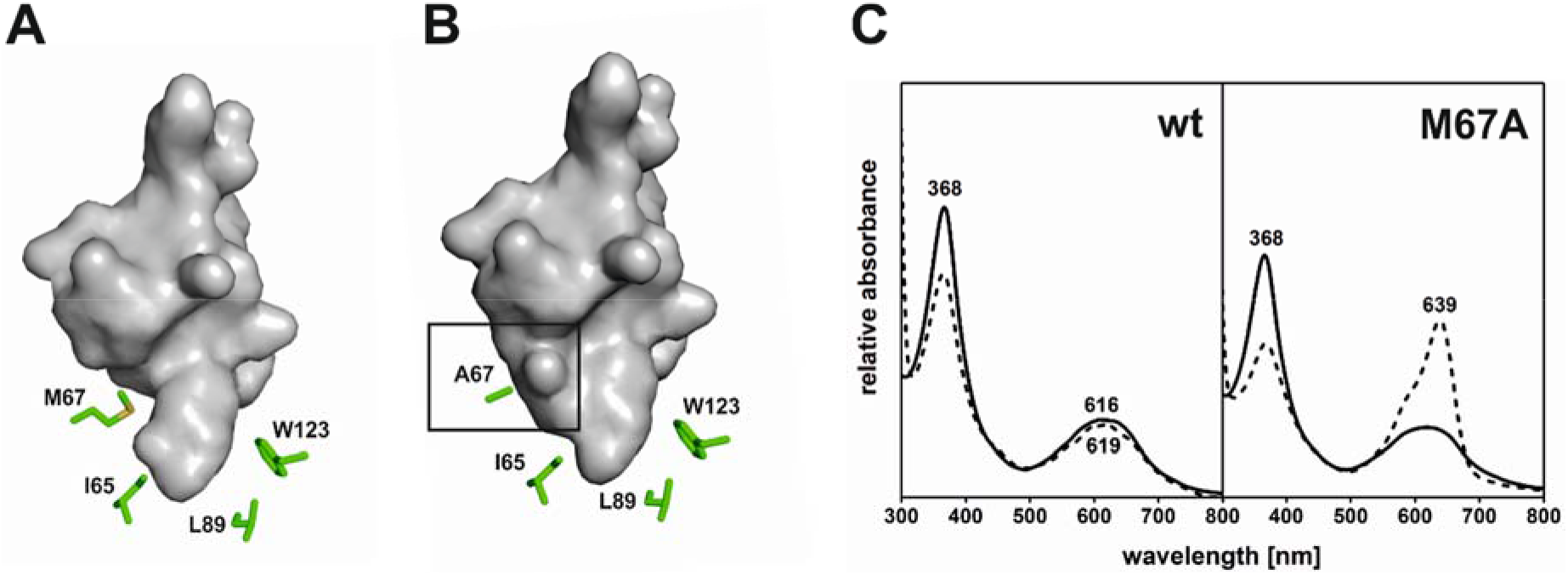
Transformation of PEB specific lyase into universal lyase. Comparison of a cavity model of the phycobilin binding pocket of *Gt*CPES (**A**) and *Gt*CPES_M67A (**B**). Model was generated using Pymol with aid of surface cavity mode (35). **C.** Absorption spectra of free 3(*E*)-PCB (solid line) compared with absorption spectra after addition of wt *Gt*CPES (left panel) and *Gt*CPES_M67A (right panel) (dashed line). Absorbance maxima are indicated.

**TABLE 3.** *In vitro* binding and transfer to *Pm*CpeB of PEB and PCB after incubation with different *Gt*CPES variants. Melting points (in °C) of proteins determined in triplicate and absorption/emission maxima of binding/transfer are presented. (Exc. 550 nm PEB; Exc. 600/620 nm PCB)

| *GtCPES*<br>(variant) | $T_m$<br>[°C] | 3(Z)-PEB | | | | | 3(E)-PCB | | | | |
| --- | --- | --- | --- | --- | --- | --- | --- | --- | --- | --- | --- |
| | | Binding | $A_{max}$<br>[nm] | $E_{max}$<br>[nm] | Transfer | $E_{max}$<br>[nm] | Binding | $A_{max}$<br>[nm] | $E_{max}$<br>[nm] | Transfer | $E_{max}$<br>[nm] |
| I65A | 47.5 ± 0.5 | yes | 555,<br>599 | --- | yes | 568 | no | 616 | --- | no | 640 |
| M67A | 53.3 ± 0.3 | yes | 553,<br>598 | 635<sup>b</sup> | yes | 569 | yes | 639 | 662 | yes | 651 |
| M67V | 52.3 ± 0.4 | yes | 555,<br>597 | --- | yes | 567 | yes | 638 | 663 | no | 640 |

*a*, Absorbance spectrum differs from wt*GtCPES*:bilin concerning extinction. *b*, fluorescence only detected for *in vivo* binding. ND means not determined, n.d., not detectable.

**FIGURE 6.**
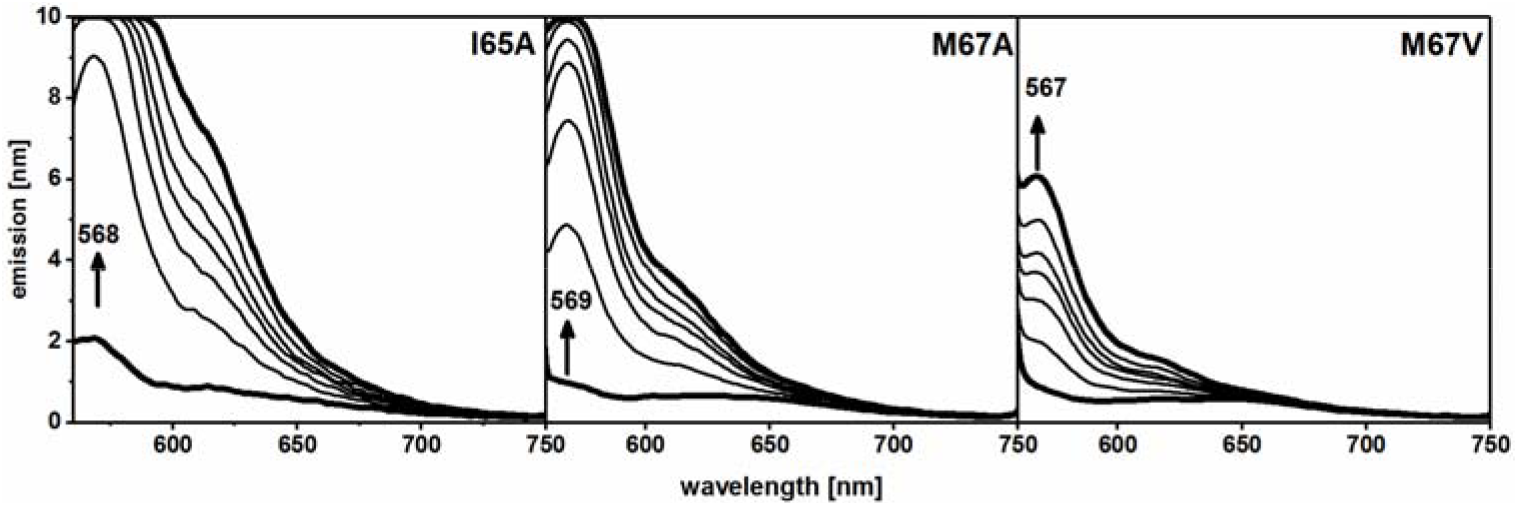
Transfer of 3(*Z*)-PEB to *Pm*CpeB by *Gt*CPES variants with larger binding pocket. Apo-*Pm*CpeB was incubated with *Gt*CPES variants preincubated with PEB. Emission spectra (λ_ex_ = 550 nm) at 1, 5, 10, 15, 20, 30, and 45 min after addition of *Pm*CpeB were taken. First and last spectra are shown by bold lines. Emission maxima are given and the course of fluorescence emission changes is indicated by arrows.

Interestingly, both methionine variants were also able to bind PCB. Incubation with PCB resulted in a shift of the PCB absorption maximum to higher wavelengths (639 nm) together with an increase of extinction reflecting phycobilin binding (Figure 5). PCB binding furthermore also resulted in a fluorescent complex formation with an emission maximum at 662 nm (Table 3). As both methionine variants displayed the same spectroscopic behavior, all following experiments were only conducted employing the M67A variant.

### GtCPES_M67A binds PCB with high affinity in vitro and in vivo

With respect to the gained PCB binding ability of the M67A variant, we next wanted to determine the affinity towards PCB. One way to do so is to investigate whether the CPES variant will form a complex with PCB when all necessary genes are coexpressed in *E. coli* (14,24) Coexpression resulted in intensely colored cells for *Gt*CPES_M67A with PEB as well as PCB (Figure 7). This complex remained stable through the affinity chromatography indicating strong binding of PCB. The latter was confirmed by determining the binding affinities for 3(*Z*)-PEB and 3(*E*)-PCB. The calculated K_D_ values showed strong binding with 0.044 μM and 7 μM for PEB and PCB, respectively (Table 2). Wild type *Gt*CPES on the other hand only formed an *in vivo* complex with PEB but not PCB ((14) and Figure 7). During the subsequent purification process of *Gt*CPES, the phycobilin remained bound to M67A as monitored by UV/vis-spectroscopy (Figure 7B). Absorption spectra of the elution fraction from affinity chromatography showed binding spectra with maxima similar to *in vitro* binding of 3(*E*)-PCB and 3(*Z*)-PEB by *Gt*CPES_M67A (Table 3). Moreover, binding of PCB resulted in the formation of a fluorescent complex with *Gt*CPES_M67A. This is in agreement with the complex formed *in vivo* (Table 3). Extinction coefficients for formed complexes with *Gt*CPES_M67A were determined assuming that all PBP-lyase molecules were loaded with PCB or PEB. Resulting coefficients are 29.56 mM^−1^cm^−1^ (*Gt*CPES_M67A:PEB) at 600 nm and 15.23 mM^−1^cm^−1^ (*Gt*CPES_M67A:PCB) at 640 nm in sodium phosphate buffer. Additionally, *in vivo* formed *Gt*CPES_M67A:PEB complex was also able to transfer loaded PEB to *Pm*CpeB detected by fluorescence spectroscopy (data not shown). Here, emission maxima were similar to *in vitro* formed complex mediated transfer.

**FIGURE 7.**
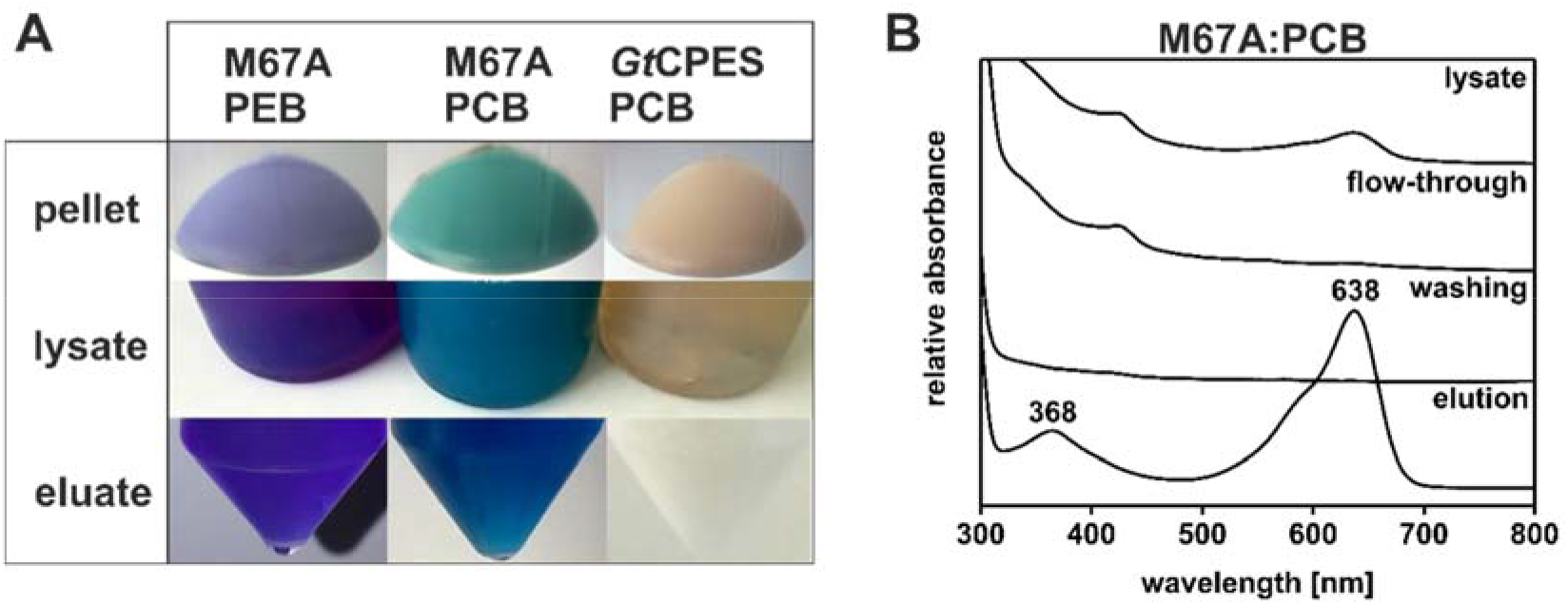
*In vivo* complex formation of *Gt*CPES_M67A with PCB and PEB. Coproduction of wt*Gt*CPES and variant *Gt*CPES_M67A with synthesis enzymes for PCB (Ho1, PcyA) and PEB (Ho1, PebS) were conducted in *E. coli* BL21(DE3). Formed *Gt*CPES_M67A:bilin complexes were purified by affinity chromatography and detected by color formation (**A**) and UV/vis-spectroscopy (**B**; *Gt*CPES_M67A:PCB). Absorption maxima in elution fraction are labelled. Coproduction of wt*Gt*CPES, Ho1 and PcyA shows no color (control).

### GtCPES_M67A transfers PCB to Cys^82^ of PmCpeB but not to CpcB

Subsequently, we were interested whether the *Gt*CPES variant M67A exhibited the ability to mediate the transfer of 3(*E*)-PCB to Cys^82^ of *Pm*CpeB. At first, spontaneous attachment of 3(*E*)-PCB was monitored by fluorescence spectroscopy (Figure 8A). Here, fluorescence spectra revealed slow but continuous emission increase over time with maximum wavelengths of 640 nm (*Pm*CpeB), 642 nm (*Pm*CpeB_C82A) corresponding to unspecific interaction of PCB with the apo-PBP. Lyase mediated product formation was detected after adding PBP subunit (*Pm*CpeB, *Pm*CpeB_C82A) to *in vitro* formed complex of *Gt*CPES_M67A and 3(*E*)-PCB, as described before. All spectra differed from spontaneous attachment with the emission intensity of the first spectrum after 1 min reaction time representing the fluorescent complex between 3(*E*)-PCB and *Gt*CPES_M67A with maximum around 663 nm (Figure 8D and Table 2). Over time, this emission maximum shifted to shorter maxima wavelengths (651 nm) indication transfer of PCB to the apo-protein. Similar fluorescence emission maxima (E_max_ =647 nm) were observed using CpcS lyase mediated PCB transfer to the CpcB subunit (Figure S2). The control experiment employing CpeB_C82A did not show any changes in absorption or fluorescence and only displayed the specific fluorescence background of *Gt*CPES:PCB (Figure 8B,E). Covalent attachment of PCB to the apo-CpeB was further confirmed using zinc blot analysis (data not shown) (25). Unfortunately, attempts to detect chromopeptides using mass spectrometry failed.

**FIGURE 8.**
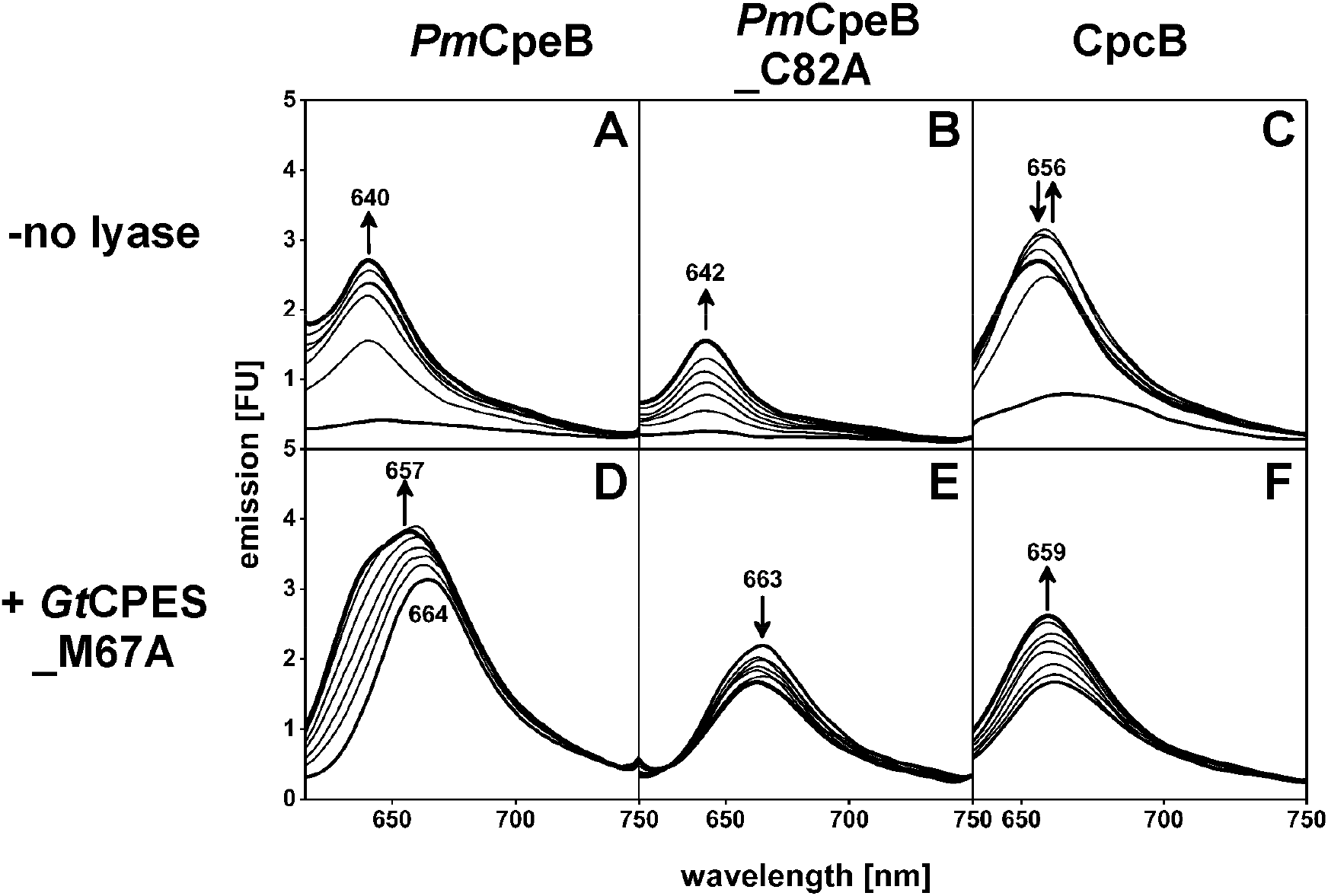
Spontaneous and *Gt*CPES_M67A mediated PCB chromophorylation of *Pm*CpeB, *Pm*CpeB_C82A and CpcB. The apo-PBPs *Pm*CpeB, *Pm*CpeB_C82A and CpcB were incubated with PCB (- no lyase) or *Gt*CPES_M67A:PCB complexes. Emission spectra (λ_ex_ = 600 nm (**A,B,D,E**); λ_ex_ = 620 nm (**C and F**)) of 1, 5, 10, 15, 20, 30, and 45 min after addition of apo-PBPs were detected. First and last spectra are shown by bold lines. Emission maxima are given and the course of fluorescence emission changes is indicated by arrows.

In a final experiment, we asked the question whether the PCB-binding *Gt*CPES_M67A variant would be able to transfer PCB to the phycocyanin β-subunit (CpcB) of *Synechococcus* sp. PCC7002. Unfortunately, here the results were not conclusive. The presence of the lyase led to an attachment product that showed fluorescence emission at 659 nm, only ~3-4 nm different to the spontaneous attachment (Figure 8C,F). However, the emission maximum also differed significantly from that mediated by the CpcS lyase, which resulted in a CpcB with a fluorescence emission at 647 nm (Figure S3). We conclude that *Gt*CPES_M67A is not able to quantitatively transfer PCB to CpcB. Whether small amounts of PCB were correctly transferred to CpcB has to be determined.

## DISCUSSION

We have previously initially characterized and crystallized the PBP lyase CPES from *G. theta*. PBP lyases are proteins that assist the correct attachment of light harvesting phycobilin pigments to conserved cysteine residues within a phycobiliprotein (15). Once all phycobilins are attached, the individual phycobiliproteins assemble into larger structures, either into phycobilisomes (cyanobacteria and rhodophytes) or into soluble heterodimers (cryptophytes). *Gt*CPES is an S-type PBP lyase and was postulated to attach PEB to Cys^82^ of the β-subunit CpeB. Although we were not able to show the transfer to the PBP subunit from *G. theta*, a transfer was observed using CpeB from the cyanobacterium *Prochlorococcus marinus* MED4. Both proteins share 55% sequence homology. We therefore conclude that *Gt*CPES is a highly specific lyase for the attachment of 3(Z)-PEB also to β-Cys^82^ of CpeB of *G. theta*. This β-subunit has in total three PEB molecules bound: an additional one with a double linkage at Cys^50,61^ and one at Cys^155^ (6,7). Accordingly, it is reasonable that *Gt*CPES is highly specific and is only able to transfer 3(Z)-PEB but not 15,16-DHBV, although binding of both bilins is observed. *Gt*CPES shows a very narrow substrate specificity to the phycobilin it is transferring. This is mainly due to a very confined binding pocket, which is characterized by several bulky amino acid residues that restrict binding to bilins containing a C15-C16 double bond like PCB or phytochromobilin (PФB) (14). However, S-type lyases from certain cyanobacteria that only possess the phycobilin PCB, i.e. CpcS, appear to have a broader substrate specificity. The CpcS lyase from *Thermosynechococcus elongatus* for instance is an universal lyase being able to bind and transfer PEB, PCB and PФB (23). Within this current study, we identified several amino acid residues that are important for phycobilin binding, transfer and substrate specificity. PEB binding is primarily mediated by two glutamate residues, E136 and E168. Both residues are facing the interior of the pocket and are likely involved in coordinating two of the pyrrole nitrogens of the bilin. E136 is highly conserved in the family of S-type lyases. W75 is also highly conserved within the S-type lyases and located at the upper rim of the barrel. Here, an involvement in the interaction with the apo-PBP is postulated. Interestingly, the *Gt*CPES_W75A variant had a significantly higher binding affinity to the substrate. We therefore initially postulated that a high affinity retains the bilin within the lyase and therefore prevents the transfer to the apo-protein. However, this is in contrast to the data observed for the *Gt*CPES_M67A variant that displayed an even higher binding affinity towards PEB (0.044 μM for M67A vs. 0.86 μM for W75A) implying that the affinity to the substrate does not determine whether the bilin is transferred or not. We rather postulate that a correct interaction with the target PBP must be present for a correct transfer to occur. This postulate is in conclusion with our data showing that the *Gt*CPES_M67A variant is unable to transfer the bound PCB to the CpcB subunit, but only to the CpeB subunit. There might be smaller structural differences that determine the specific interaction of the CPES lyase with its corresponding (or in our case homolog) apo-protein.

Finally, we identified a crucial amino acid residue for substrate specificity. M67 determines the narrow substrate specificity of PEB-specific S-type lyases. This residue is highly conserved among CpeS lyases and is only substituted by an isoleucine in some cases (Figure S1). Within the PCB specific S-type lyases, this position is occupied by valine. With its shorter side chain, a valine residue at this position widens the pocket and allowing the more rigid PCB to enter the pocket. The same hold true for our M67A and M67V variants. When the long side chain of methionine is modified to either alanine or valine, the substrate specificity is broadened and the protein can bind both, PEB and PCB. We therefore hypothesize a crucial function for amino acid residue at position 67 in determining the substrate specificity of S-type lyases. Overall, our data provide very strong support for a dual function of the lyase both as bilin-selective binder and bilin-prepositioning chaperon for the hand-over to a dedicated target.

## Experimental procedures

### Materials

All chemicals were American Chemical Society grade or better unless specified otherwise. Expression vector pCOLADuet-1 was obtained from Novagen; pASK-IBA7+ was from IBA Life Sciences, pET-Duet1 from Merck KgaA, and pGro7 was from TaKaRa. Strep-Tactin®-Sepharose from IBA, and TALON® metal affinity resin from Clontech were used. HPLC-grade acetone, acetonitrile, formic acid, and spectroanalytical grade glycerol were obtained from J.T. Baker. Sep-Pak cartridges were obtained from Waters.

### Construction of expression plasmids and site directed mutagenesis

For construction of pCOLA*cpeB* a synthetic gene codon optimized for *E. coli* K12 (GENEius algorithm, MWG Eurofins Operon) encoding the *cpeB* gene from *Prochlorococcus marinus* MED4 was PCR amplified with primers (Table 4) encompassing selected recognition sites (*Eco*RI, *Hind*III) for cloning into pCOLADuet (Novagen).

**TABLE 4.**
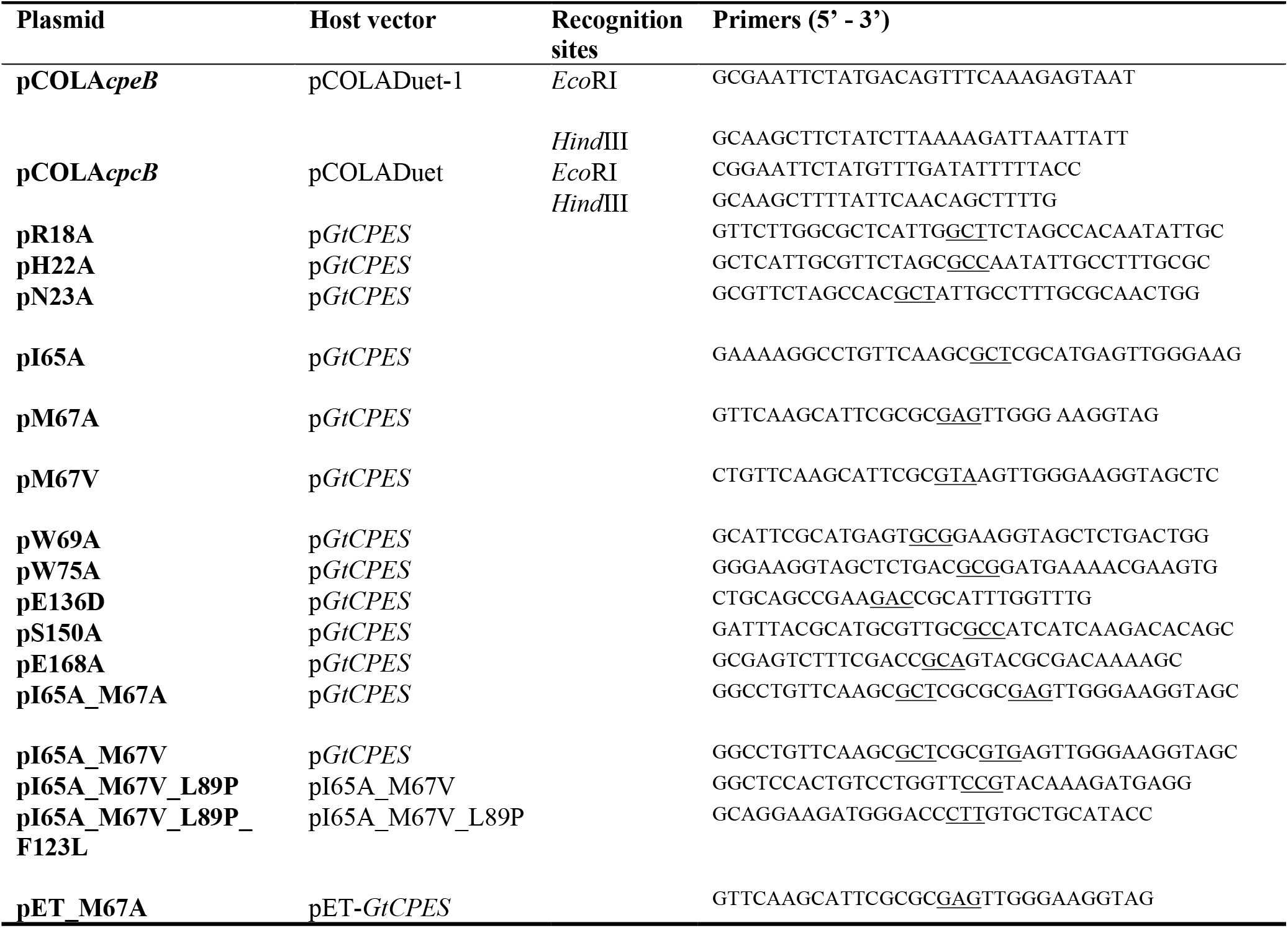
Constructed expression plasmids. Forward primers for site-directed mutagenesis are shown, reverse primer is the complement. Changed codons are underlined.

For construction of pCOLA*cpcB* the plasmid pBS150v_7002*cpcB* was used (26) This construct was a gift of W. M. Schluchter (University of New Orleans). The *cpcB* gene from *Synechococcus* sp. PCC7002 was PCR amplified (Table 4) with same recognition sites as for *cpeB* for following cloning into pColaDuet.

The construction of the plasmids p*GtCPES*, pE136A, pR146A, pR148A, pC149A, pET*GtCPES*, pTD*ho1pcyA* and pTD*ho1pebS* were described before (14,24,27) All additional site-directed variants of *Gt*CPES and *Pm*CpeB were generated from p*GtCPES* or pCOLA*CpeB* using the QuikChange^®^ site-directed mutagenesis kit (Stratagene) with help of primers listed in Table 4 (only the forward primer is shown, the reverse primer is the complement, codon changes are underlined). The resulting plasmids were verified by sequencing.

### Production and purification of recombinant proteins

*Gt*CPES and all variants as well as CpcS were produced and purified as described before (14,28) *Pm*CpeB and CpcB were produced expressing pCOLA*cpeB*, pCOLA*cpeB*_C82A or pCOLA*cpcB* in *E. coli* BL21(DE3)-RIL. Cells were grown at 37 °C in LB medium supplemented with 50 μg/ml kanamycin, and for *cpeB* with 100 mM D-sorbitol and 2.5 mM betaine to OD_578 nm_ of 0.5 - 0.6. Subsequently, cells were induced by isopropyl-β-D-thiogalactopyranoside (IPTG; 0.5 mM) and incubated overnight at 17 °C. Cells were harvested by centrifugation, resuspended in lysis buffer (50 mM Tris-HCl pH 7.5, 100 mM NaCl, 10% Glycerol) and lysed by two or three passages through LM10 Microfluidizer® High Shear fluid homogenizer (Microfluidics) (*cpeB*, *cpcB, gtCPES* constructs) at 19.000 psi or by three times 3 min sonification (50%, KE76, Sonopuls HD6600, Bandelin) (only *gtCPES* constructs).

*Pm*CpeB, its variant *Pm*CpeB_C82A and CpcB were purified by affinity chromatography using TALON® metal affinity resin (Clontech) and purification was carried out according to the manufacturer’s instructions based on sodium phosphate buffer (60 mM, 300 mM NaCl, pH 7.5). For imidazole removing a dialysis against 100fold volume of sodium phosphate buffer (60 mM sodium phosphate, 300 mM NaCl, pH 7.5) over night at 4 °C (120 rpm) was performed.

### Determination of protein concentration

Purified proteins were concentrated using Amicon® Ultra-4 Ultracel®-10K (MWCO: 10.000 Da; Merck). Protein concentrations were quantified using the calculated molar extinction coefficient ε_280_ (29).

### Coproduction of phycobilins and PBP-lyases

Heterologous coproduction of *Gt*CPES or variant *Gt*CPES_M67A and PEB biosynthesis enzymes was performed in *E. coli* BL21(DE3) containing pET-constructs of *gtcpeS* and *gtcpes_M67A* (*pET_gtCPES; pET_M67A*) and p*ho1pebS* or *pTDho1pcyA*. Cultures were grown in LB medium supplemented with 100 mM sorbitol and 2.5 mM betaine at 37 °C, 100 rpm to an OD_*578nm*_ of 0.6 prior to induction with 0.1 mM IPTG and incubated overnight at 17 °C (~16 h). After cell harvesting by centrifugation, cells were resuspended in lysis buffer (60 mM sodium phosphate, 300 mM NaCl, pH 7.5), and disrupted by pressure based homogenizer (LM10 Microfluidizer®, Microfluidics), preferably, or sonification. Purification process was described before (14)

### Determination of extinction coefficient for PBP-lyase:phycobilin complex

Extinction coefficients were determined for *in vivo* produced and purified PBP lyase:phycobilin-complexes. Here, 30 μM of PEB or PCB were added to 20 μM of *Gt*CPES or variant *Gt*CPES_M67A, total volume was 550 μl. Phycobilin excess was removed by gel filtration (PD Minitrap G25 Sephadex column, GE healthcare) and performed according to the manufacturer’s instructions with spin protocol based upon centrifugation (3000 rpm, 2 min). PBP-lyase concentration was determined by UV/vis-spectroscopy between all steps. Assuming that all PBP-lyase molecules were loaded with phycobilins, extinction coefficient for PBP-lyase: PEB complex at 600 nm and PBP-lyase:PCB complex at 640 nm were calculated with aid of Lambert-Beer law.

Determination of extinction coefficient for that complex (ε_600_ = 27.79 mM^−1^ cm^−1^) enabled the calculation of complex bound protein concentration under the assumption that all *Gt*CPES molecules were present in complex with PEB.

### Thermal shift experiments

The temperature of the unfolding transition midpoint, for excluding effect due to denaturation of *Gt*CPES variants, was determined in thermal shift experiments (30,31) For determination fluorescent protein stain SYPRO® Orange (5000x, Sigma-Aldrich) and CFX Connect Real-Time PCR Detection system (Bio-Rad) were used. Microplates were obtained from Biozym. With exclusion of light 50 μl total volume composed of 5 μl protein (2 mg ml^−1^), 5 μl SYPRO® Orange (100x) and sodium phosphate buffer (60 mM sodium phosphate, 300 mM NaCl, pH 7.5) were tested in triplicate. Controls were conducted without protein component. Melting temperatures were determined automatically by software CFX Maestro.

### Preparation of phycobilins

PCB (3(*E*)-PCB) was isolated from *Spirulina* cells as described previously (32). Production, purification and preparation of PEB were described before (14) but PEB was concentrated by Advance Alpha 2–4 LSCplus freeze dryer (Christ). The vacuum was set to 0.04 mbar with ice condenser set at −40 °C. Phycobilins were resuspended in an appropriate amount of DMSO before use. The isolated phycobilins were analyzed in terms of isomers via HPLC using Luna 5μ C18 column (Phenomenex) and stored at −20 °C on silicagel orange in the dark. If isomer ratio showed strong excess of *E* or *Z*-isomer for PEB or PCB, concentrations were determined using ε_571_: 46.9 mM^−1^cm^−1^ (3(*E*)-PEB) in MeOH/5% HCl and ε_685_: 37.15 mM^−1^cm^−1^ (3(*E*)-PCB) in MeOH/2.5% HCl (33). Due to the absence of a reported extinction coefficient for 3(*Z*)-PEB ε_571_ of the related 3(*E*)-PEB was used. 3(*Z*)-PEB was directly applied in experiments without separation of isomers and further purification because of *Z*-isomer excess.

### Phycobilin binding and transfer by *Gt*CPES variants

For binding studies excess of *Gt*CPES or *Gt*CPES variant (20 μM), respectively, and phycobilin (5 μM) were mixed in sodium phosphate buffer, pH 7.5. Absorption spectra (Agilent 8453 UV-visible spectrophotometer) and in case of transfer studies fluorescence emission spectra (Series 2, FA-256, Aminco Bowman) of the mixture and free phycobilins in the same buffer were detected. Measurements were done at 60% sensitivity and 975 V. For transfer studies 20 μM *Pm*CpeB, *Pm*CpeB_C82A or CpcB was added 2 min after *Gt*CPES (variant):phycobilin-complex formation and fluorescence emission was measured after 1, 5, 10, 15, 20, 30 and 45 min (Exc. 550 nm PEB; Exc. 600/620 nm PCB). All samples were incubated and measured at room temperature. After an incubation time of 45 min all sampled were prepared for and separated by SDS-PAGE on a 12.5% gel. Proteins were transferred to PVDF membrane that was subsequently incubated in 1.3 M zinc acetate for 1 h at 4 °C, and afterward zinc-enhanced fluorescence of proteins covalently associated with bilins was visualized under UV light (312 nm) (25)

### Spectroscopic analysis of phycobilin binding

Increasing amounts of *Gt*CPES (variant) (2-15 μM) were added to 10 μM (final concentration) 3(*Z*)-PEB or 3(*E*)-PCB in a final volume of 200 μl of sodium phosphate buffer, pH 7.5, under exclusion from light. After incubation for 2 min absorbance spectra were recorded using Agilent 8453 UV-visible spectrophotometer. The method was described by Frankenberg and Lagarias (34) and was adapted, here. Analysis of spectra was performed using Microsoft Excel.

To obtain phycobiliprotein lyase dissociation constants, absorbance differences (Δ*A*) at the λ_max_ of each complex were plotted as a function of *Gt*CPES concentration with aid of Microsoft Excel. Dissociation constants K_D_ were obtained by fitting the parameters of the equation (1) to the experimental data, where [P_total_] describes the total concentration of *Gt*CPES, [L_total_] of phycobilin and [PL] is the concentration of complex.

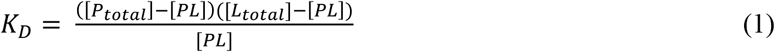

### Data and image processing

Spectra were generated in Origin® data analysis/graphic software, models of protein structure or ligand binding pocket were edited with aid of published *Gt*CPES crystal structure (PDB code 4TQ2) by Pymol (35). Cavity model of *Gt*CPES binding pocket was displayed by surface cavity mode of Pymol.

## ACKNOWLEDGEMENTS

This work was supported by a grant from the Deutsche Forschungsgemeinschaft to NFD. We like to thanks Wendy Schluchter and Kai-Hong Zhao for the gift of the expression plasmids.

## AUTHORS CONTRIBUTION

NT, KEO, AP, EH and NFD designed the research, NT, KEO, MA performed the experiments, AP helped analyzing data, NT, KEO, NFD analyzed data and wrote the manuscript. All authors approved the manuscript.

