## supporting information Figures 1-3 for "Exchange of a single amino acid residue in the cryptophyte phycobiliprotein lyase *Gt*CPES expands its substrate specificity"


---

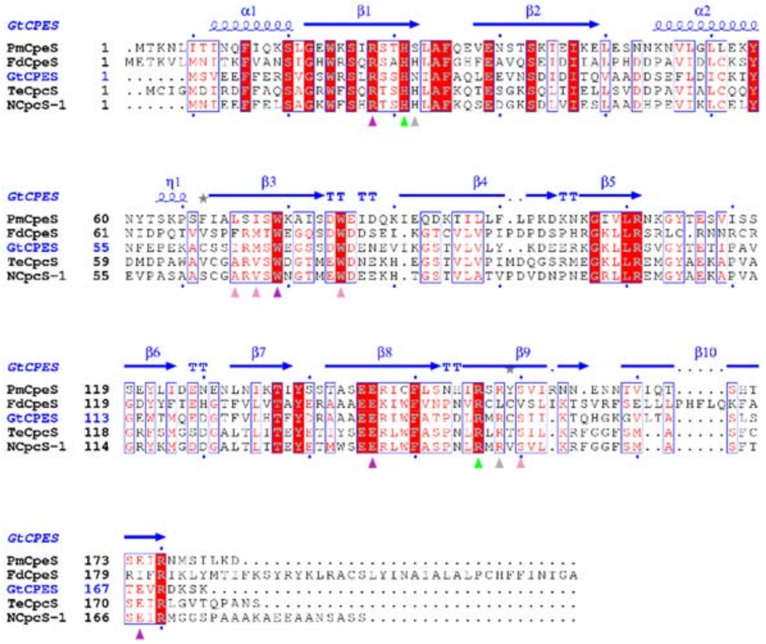


**Figure S1. Sequence alignment of different S-type lyases.** PmCpeS: *P.marinus* CpeS (Uniprot Q7V2Z2); FdCpeS: *Fremyella diplosiphon* CpeS (Uniprot A5HIW6); GtCPES: *G.theta* CPES (Uniprot L1JFC7); TeCpcS: *T.elongatus* CpcS (Uniprot Q8DI9I); NCpcS-1: *Nostoc* sp. PCC7120 CpcS 1 (Uniprot Q8YZ70). The secondary structure of *Gt*CPES (PDB code 4TQ2) is shown in blue. Residues discussed in the main text are marked with triangels with color coding equivalent to Figure 2, representing the effect of a site-specific exchange on *Gt*CPES function: white – no change; green – structural defect; purple – bilin binding; pink – bilin transfer and selectivity. Alignment generated with ClustalΩ as implemented in the EBI-Tools (1,2). Figure generated with ESPript (http://espript.ibcp.fr, (3))

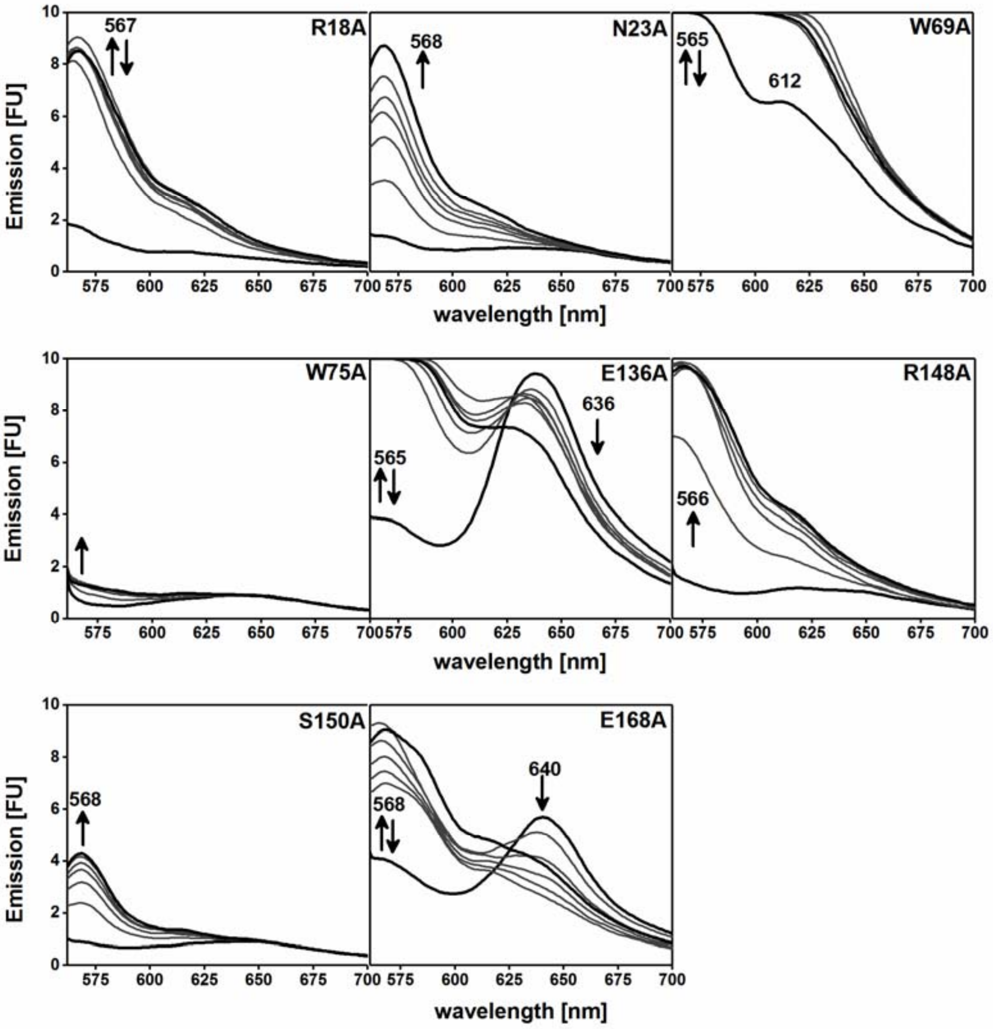


**Figure S2. Transfer of 3(*Z*)-PEB to *Pm*CpeB by *Gt*CPES variants.** The apo-PBP *Pm*CpeB was incubated with PEB and *Gt*CPES variants (**A-K**). Emission spectra ($\lambda_{ex} = 550$ nm) of 1, 5, 10, 15, 20, 30, and 45 min after addition of *Pm*CpeB, respectively, were detected. First and last spectra are shown by bold lines. Emission maxima are given and the course of fluorescence emission changes is indicated by arrows. Please note that spectra shown for R18A, N22A, W69A, W75A, R148A, S150A and E168A are taken from Figure 4 and only shown here for completeness.

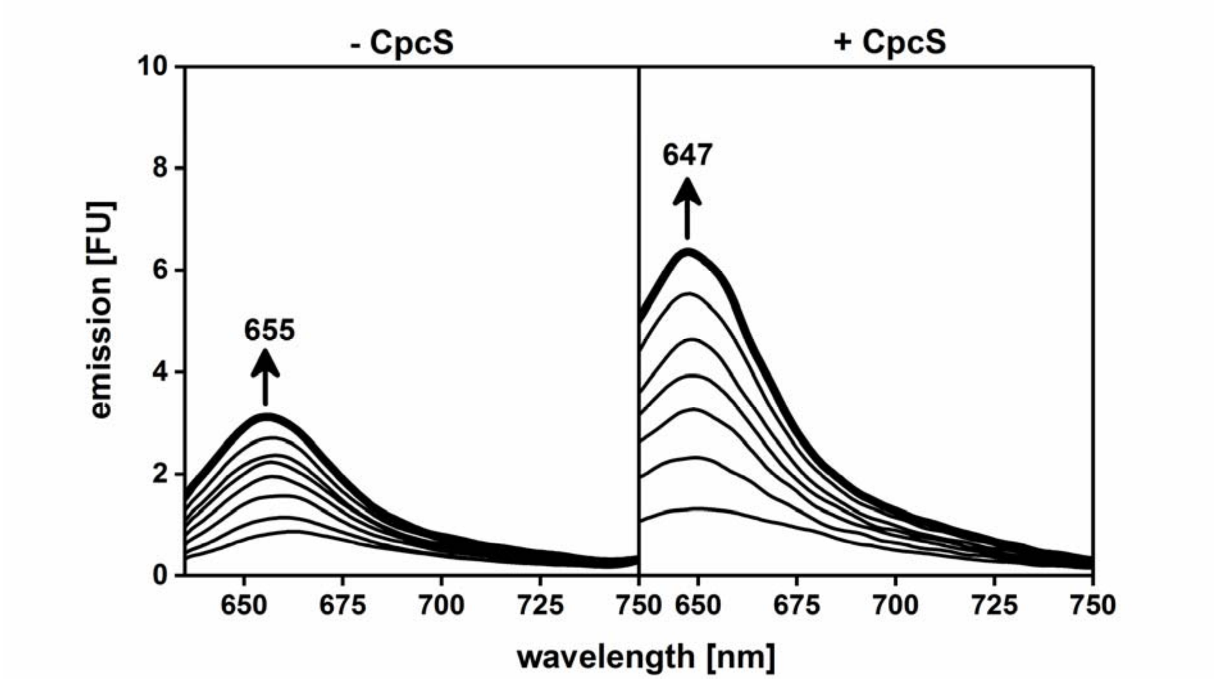


**Figure S3. Transfer of 3(*E*)-PCB to *Synechococcus* CpcB by *Nostoc* CpcS.** The apo-PBP CpcB was incubated with either 3(*E*)-PEB istself (-CpcS) or with CpcS preincubated with submolar concentration of 3(*E*)-PCB. Emission spectra ($\lambda_{ex}$ = 620 nm) of 1, 5, 10, 15, 20, 30, and 45 min after addition of CpcB were detected. First and last spectra are shown by bold lines. Emission maxima are given and the course of fluorescence emission changes is indicated by arrows.
